# IGF2BP3 is essential for the growth heterogeneity of colorectal adenoma cells by regulating MYC

**DOI:** 10.1101/2025.09.08.674833

**Authors:** Tomohiko Sunami, Roberto Coppo, Kunishige Onuma, Jumpei Kondo, Hiroyuki Uematsu, Daisuke Hoshi, Yoshihiro Yamamoto, Osamu Kikuchi, Shinya Ohashi, Yasuhide Takeuchi, Kazuto Kugou, Yoshinori Hasegawa, Yoshiro Itatani, Kazutaka Obama, Yoshitaka Hippo, Manabu Muto, Atsushi Yamada, Masahiro Inoue

**Author notes:** Current affiliation of J.K.: Division of Health Sciences, Department of Molecular Biology and Clinical Investigation, Graduate School of Medicine, The University of Osaka, Japan; D.H.: Department of Oncologic Pathology, Kanazawa Medical University, 1-1 Daigaku, Uchinada, Kahoku, Ishikawa 920-0293, Japan. Correspondence: Masahiro Inoue Graduate School of Medicine Kyoto University, Department of Clinical Bio-resource Research and Development, Med-Pharm Collaboration Bldg 503, Shimoadachi-cho 46, Sakyou-ku, Kyoto, 606-8304, Japan, Atsushi Yamada Department of Medical Oncology, Kyoto University Graduate School of Medicine, 54 Shogoin Kawahara–cho, Sakyo–ku, Kyoto 606–8507, Japan.

## Abstract

Adenoma is a major precancerous lesion in colorectal cancer (CRC), and the adenoma-carcinoma sequence is a well-known multistep progression to CRC caused by the accumulation of genetic mutations. On the other hand, the non-genetic mechanisms in the adenoma-carcinoma sequence remain largely unknown. In this study, organoids were established from 31 colorectal adenomas from 19 patients. Some adenomas showed heterogeneity in the proliferative potential of single cells, and this heterogeneity was regulated by non-genetic mechanisms. *IGF2BP3* was identified as a differentially expressed gene between organoids with different growth patterns. IGF2BP3 positively regulated MYC expression at the transcriptional level and negatively regulated it at the translational level. This promoted high proliferative potential with high levels of oxidative phosphorylation, while allowing cells to avoid MYC-induced cell death. IGF2BP3 affected the tumorigenicity of mouse adenomas *in vivo*. Intra-tumor heterogeneity in growth potential is acquired at the precancerous stage in the adenoma-carcinoma sequence of colorectal carcinogenesis, and IGF2BP3 plays an important role in regulating MYC levels. These findings provide new insights into the non-genetic regulation of adenomas during CRC development.

## Introduction

Colorectal cancer (CRC) is one of the most common causes of cancer-related deaths globally; therefore, its prevention and early detection are important(Siegel *et al*, 2023). The adenoma-carcinoma sequence explains the multistep progression from colorectal adenoma to CRC, accompanied by the accumulation of genetic mutations(Kinzler & Vogelstein, 1996; Vogelstein *et al*, 1988). This process usually begins with a mutation in the *APC* gene, followed by further mutations in tumor suppressor genes and oncogenes, such as *KRAS* and *TP53*. There are reports that approximately 70% of CRCs are caused by this sequence(Kinzler & Vogelstein, 1996). Thus, colorectal adenomas play an important role as precancerous lesions of CRC. Accumulating evidence indicates that CRC exhibits significant intra-tumoral heterogeneity caused not only by genetic mutations, but also by epi-genetic mechanisms, such as DNA methylation, histone modification, and chromatin accessibility(Laisne *et al*, 2025). These epi-genetic processes play crucial roles in generating phenotypic diversity and cellular plasticity, which drive the dynamic and adaptive behavior of cancer cells(Househam *et al*, 2022; Tape, 2024); however, the role of epigenetic changes in adenomas is largely unknown.

In research on colorectal adenomas, a limited number of cell lines(Paraskeva *et al*, 1989) and mouse models(Moser *et al*, 1992; Shibata *et al*, 1997) have been used. In recent years, organoid culture methods have made it possible to culture cells derived from colorectal adenomas in patients, and it is now possible to assess heterogeneity within adenomas and between patients. We recently demonstrated that patient-derived CRC organoids consist of heterogeneous cells with distinct fates and plasticities during their proliferation(Coppo *et al*, 2023a). We developed a single-cell-derived spheroid-forming and growth (SSFG) assay to precisely monitor the growth fate of single cells(Coppo *et al*., 2023a; Coppo *et al*, 2023b). In these experiments, CRC cells exhibited heterogeneous proliferative properties, consisting of cells capable of forming small spheroids (S-cells) and cells generating large spheroids (L-cells). S-cells are characterized by their slow-growing capacity. In subsequent SSFG assays, they consistently formed only small spheroids. This phenotype is known as the “S-pattern”. In contrast, L-cells had a high proliferative capacity and formed both small and large spheroids. This phenotype is known as the “D (dual)-pattern”. S-cells and L-cells exhibit cellular plasticity, transitioning from the “S-pattern” to the “D-pattern”, and vice versa, under specific conditions. However, there are still important questions to be answered, such as when and how this heterogeneity arises, particularly during the transition from normal to CRC cells. Elucidating these aspects at the molecular level is essential for understanding the mechanisms of cancer progression.

In this study, using an organoid culture technique, we investigated the growth pattern of colorectal adenoma cells and revealed that the heterogeneity of growth ability, which is observed in CRC, was already acquired in some adenomas. We also found that insulin-like growth factor 2 mRNA-binding protein 3 (IGF2BP3), also known as IMP-3 played a critical role in heterogeneous growth by modulating MYC expression. A deeper understanding of the process by which colorectal adenomas develop into CRC will lead to the development of new methods to prevent CRC.

## Results

### Adenoma organoids exhibited heterogeneous growth at the single-cell level

Organoids were established from 31 colorectal adenomas from 19 patients using a previously reported method(Sato *et al*, 2011) (Table S1). No serrated polyps were included. The microscopic cell morphology retained features of the original patient’s tumor, such as the nuclear-to-cytoplasmic ratio and staining for the colonic epithelial marker pan-cytokeratin. Representative data from CP9-1 cells are shown in Fig. 1A. First, the SSFG assay(Coppo *et al*., 2023a) was performed to monitor the growth rate of individual cells in organoids from adjacent morphologically normal regions. Non-neoplastic organoids of the colon, including those from patients with familial adenomatous polyposis (FAP), cannot be maintained under adenoma culture conditions and were cultured under normal colonic organoid culture conditions(Sato *et al*., 2011). Sixty percent (3/5) of the non-neoplastic organoids formed spheroids from single cells. The three cases that formed spheroids (from non-neoplastic tissue from patients with sporadic CRC, FAP, and Lynch syndrome) all showed relatively uniform growth with an S-pattern, and none showed a D-pattern (Fig. 1B). SSFG assays were performed using adenoma organoids. The frequency of spheroid formation from single cells of the adenoma organoids was 83.8% (26/31) and 57.7% (15/26) of the spheroid-forming cells showed an S-pattern similar to that of normal colonic epithelium, while 42.3% (11/26) showed a D-pattern (Fig. 1C). There was no correlation between the D/S-pattern and histological type, size, or site of origin. One case of carcinoma in an adenoma exhibited a D-pattern. The SFC was significantly higher in the D-pattern adenomas (Fig. 1D). Monitoring the spheroid growth of single cells from adenomas revealed that each cell grew over time in both D- and S-patterns (Fig. 1E, F). The reproducibility of the SSFG assay results was confirmed for two D-pattern adenomas and two S-pattern adenomas (Fig. S1). These results indicated that the D/S-pattern is a stable feature specific to each adenoma. Thus, the D-pattern observed in CRC (Coppo *et al*., 2023a), i.e. the diversity of cell growth fates within the same tumor, has already been acquired in some adenomas.

**Figure 1.**
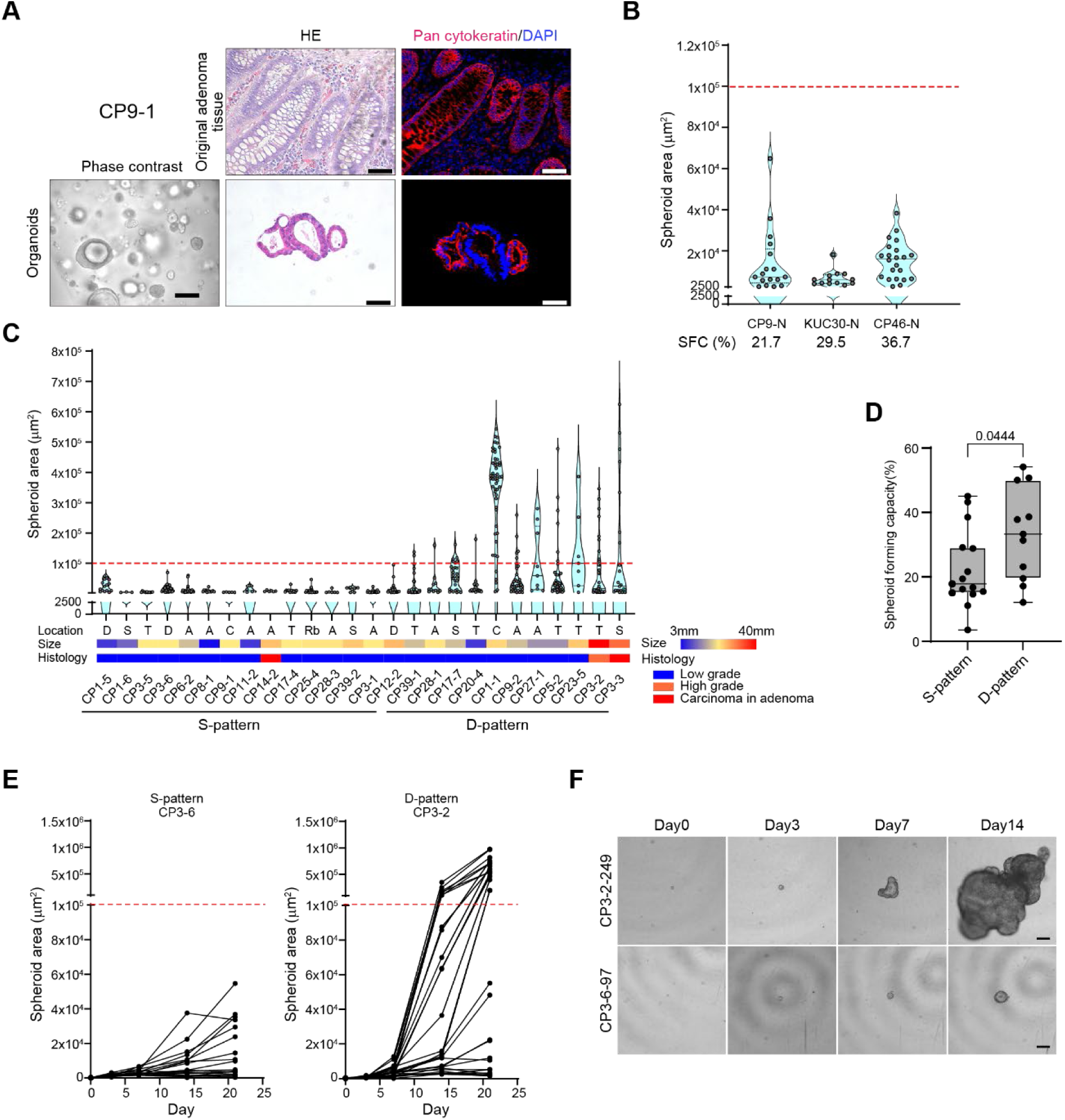
Adenoma organoids exhibited heterogeneous growth at the single-cell level **A**) Brightfield, hematoxylin and eosin (H&E), and immunofluorescent images of a representative adenoma organoid and the corresponding original adenoma tissue. Scale bar: bright field, 100 μm; others, 50 μm. **B**) Violin plots showing the results of the single-cell-derived spheroid-forming and growth (SSFG) assay for organoids from a morphologically non-neoplastic region. Each dot represents single-cell-derived spheroids. The red dashed line indicates a rounded value (1.0 × 10^5^ μm^2^) of the area at day 14. Each dot represents a growing spheroid originated from a single cell. SFC are shown below. **C**) Violin plots of the SSFG assay for adenoma organoids with size, location, and SFC indicated below. **D**) Comparison of SFC between D-/S-pattern groups. The mean ± SD is shown, tested by Fisher’s exact test. **E**, **F**) Growth curve single cells from S- and D-pattern organoids. Each line represents growth of a single cell-derived spheroids. (**E**) and phase-contrast images of the representative spheroids (**F**). Scale bar: 100 μm

### Growth heterogeneity in D-pattern organoids was epigenetically regulated

We examined whether the growth patterns were genetically determined. Because the SSFG assay monitors the growth of spheroids originating from single cells, the spheroids are clones. The clones were expanded and a second round of the SSFG assay was performed. The SFC of organoid CP1-1 in the second round was almost the same as that in the first round (Fig. 2A). One large spheroid clone (#160) exhibited a D-pattern in the second round. The other large spheroid clone (#259) showed an S-pattern by definition, but had a relatively wide growth range. Both small spheroids (#75 and #230) exhibited an S-pattern in the second round. Growth variances in large spheroid clone #259 of CP1-1, which showed an S-pattern in the second-round SSFG assay, were significantly larger than those in small spheroid clones #75 and #230 (*p* < 0.0001, F test). It should be noted that in the second-round SSFG assay, large spheroid-forming clones did not give rise to fast-growing cells, but most of them again showed a D-pattern. These results suggested that the fast-growing phenotype in the D-pattern was not genetically determined. Once isolated, all the small spheroid clones gave rise to small spheroids. These features were confirmed with another adenoma organoid (CP9-2), except for the large spheroid-forming clone #136, which exhibited an S-pattern in the second-round assay. Thus, the growth features of adenoma cells were similar to those of CRC cells(Coppo *et al*., 2023a).

**Figure 2.**
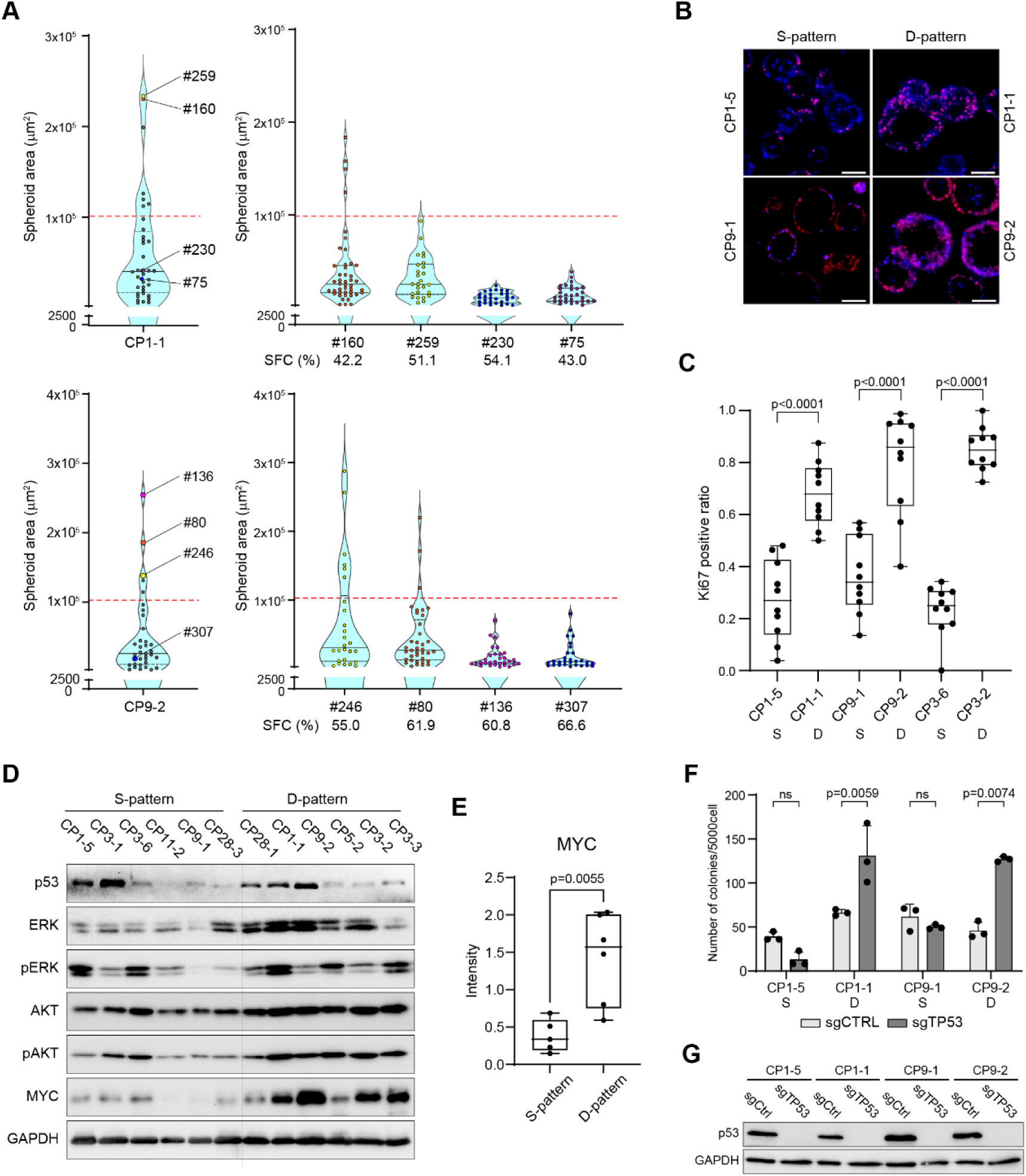
Growth heterogeneity in D-pattern organoids was epigenetically regulated **A**) Violin plots of the SSFG assay for the adenoma organoids, C1-1 and CP9-2. Four selected spheroids are indicated (left). Each clone is subjected to the next round of the SSFG assay (right). The spheroid-forming capacity (SFC) is shown below. **B**, **C**) Immunofluorescence images of Ki-67 in the D/S-pattern (**B**) and quantitative comparison of the Ki-67-positive ratio (**C**). Scale bar: 50 μm **D**) Western blotting analysis of the indicated proteins and their phosphorylated forms among individual organoids. **E**) Quantitative comparison of MYC protein expression levels in (D) between S-pattern and D-pattern organoids. **F**) Quantitative analysis of soft-agar colony-formation assay. Each dot represents an independent experiment. **G**) Western blotting analyses of p53 in organoids infected with lentiviruses expressing Cas9 and the TP53 sgRNA (sgTP53) or a non-targeting sgRNA (sgCtrl).

Next, we compared the biological characteristics of the D- and S-patterns. The Ki67-positive cell ratio was higher in the D-pattern organoids than the S-pattern organoids (Fig. 2B, C), while the cleaved-caspase-3-positive ratio was approximately the same between patterns (Fig. S2A), suggesting that the differences in spheroid growth were related to proliferation. Regarding molecules related to CRC, there were differences in the protein levels of p53 and phosphorylated ERK and AKT between individual spheroids, but there was no bias between the S- and D-patterns (Fig. 2D). In contrast, MYC expression was significantly higher in the D-pattern organoids than the S-pattern organoids (Fig. 2D, 2E).

Next, whole-exome sequencing was performed on six cases of S-pattern and D-pattern adenomas from three patients (CP1, CP3, and CP9) to investigate whether there were any differences in gene mutations between S-pattern and D-pattern adenomas. *APC* mutations were detected in all six cases, but no mutations were detected in *TP53* or *BRAF* (Fig. S2C). A *KRAS* mutation was detected in only one case, which showed an S-pattern. No mutations were found in CRC-related genes that completely distinguished between the S-pattern and D-pattern groups (Table S3). The mean mutational burden was higher in the D-pattern group; however, this difference was not statistically significant (Fig. S2B).

Next, soft-agar colony-formation assays were performed. This assay evaluates anchorage-independent growth *in vitro* and is widely used to evaluate malignant transformation. There was no difference in the rate of colony formation between the S-pattern and D-pattern adenomas (Fig. 2F). However, when *TP53* was knocked out (Fig. 2G), the rate of colony formation increased only in D-pattern adenomas. These results suggested that D-pattern adenomas are at a more advanced stage of progression in the adenoma-carcinoma sequence (Fig. 2F). Organoids with *TP53* knocked out were transplanted into mice, and none of them were tumorigenic (data not shown).

### IGF2BP3 was upregulated in D-pattern organoids

The gene expression profiles of the S-pattern (CP1-5, CP3-5, and CP9-1) and D-pattern (CP1-1, CP3-2, and CP9-2) organoids were compared using microarray analysis. GO biological processes and hallmark pathway analyses were performed to compare the D-pattern and S-pattern adenoma organoids. Multiple GO terms related to the cell cycle, mitochondrial function, and biogenesis (Fig. 3A), along with pathways related to the cell cycle, MYC, mitochondrial function, and metabolism (Fig. 3B), were upregulated and enriched in the D-pattern organoids. No mutations were detected in p53 (Fig. S2C), and no differences were observed at the protein level in the examined organoids (Fig. 2D, 2G). Pathway analysis showed that the p53 pathway was downregulated in the D-pattern organoids (Fig. 3B). In the GSEA, the signature of mitochondrial gene expression and translation was upregulated, and the p53 pathway was downregulated in the D-pattern organoids (Fig. 3C). These results suggest that D-pattern organoids have high mitochondrial activity and functional suppression of the p53 pathway.

**Figure 3.**
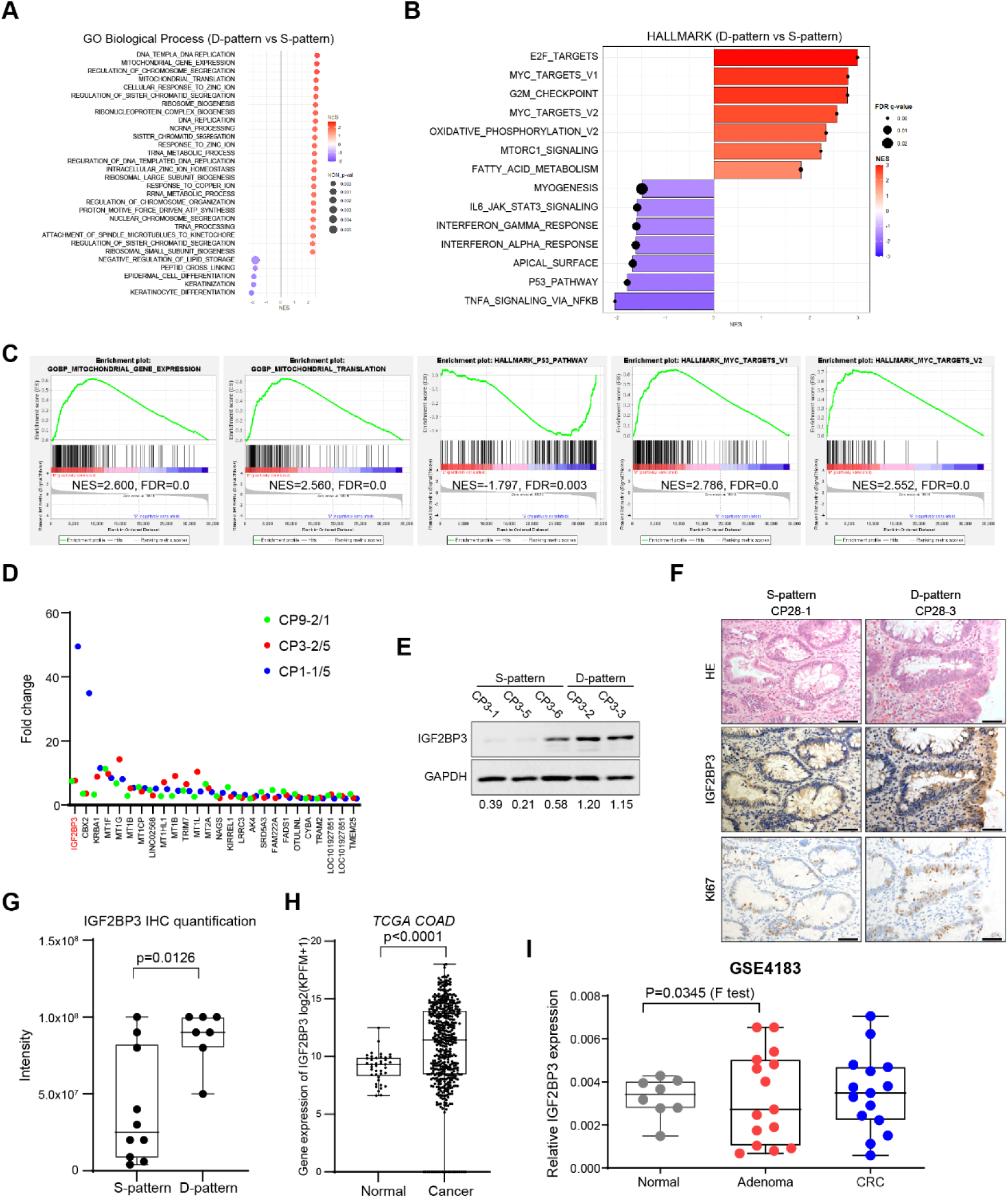
IGF2BP3 was upregulated in D-pattern organoids **A, B**) Gene Ontology (GO) analysis of biological processes (**A**) and hallmark pathways (**B**) between D/S-patterns. GO terms are shown on the y-axis, with normalized enrichment scores (NES) and *p*-values displayed on the x-axis. The bubble size is proportional to the Nominal (NOM) *p*-value and uses a color gradient for the NES in (**A**). (**A**, **B**) Red gradient colors represent upregulated processes enriched in the D-pattern, while blue gradient colors indicate downregulated processes. **C**) Gene set enrichment analysis (GSEA) of the transcriptome data using indicated signatures comparing the D/S-pattern. **D**) Fold changes of commonly upregulated genes in the D-pattern organoids. A comparison between each indicated S-pattern and D-pattern organoid is shown. **E**) Western blotting analyses of the IGF2BP3 protein in adenoma organoids. Relative expression levels were normalized to GAPDH levels and the quantified values are shown below each band. **F, G**) Representative H&E and immunohistochemical staining images of IGF2BP3 and Ki67 (**F**), and quantitative analysis of IGF2BP3 intensity comparing D/S-patterns (**G**). Scale bar: 50 mm. **H**) Gene expression of *IGF2BP3* in normal colon and colorectal cancer (CRC) from The Cancer Genome Atlas. **I**) Gene expression of *IGF2BP3* in normal colon, adenoma, and CRC from GSE4183.

Of the 25 genes that were commonly upregulated in the D-pattern organoids compared to the S-pattern organoids from the adenomas of the three patients, the gene with the largest difference in expression levels was *IGF2BP3*, an N6-methyladenosine (m6A) RNA methylation regulatory factor (Fig. 3D). The D-pattern showed higher IGF2BP3 protein expression levels than the S-pattern derived from the same patient’s adenoma (Fig.3E). Immunostaining of the original adenoma that underwent the SSFG assay showed that the D-pattern had a higher IGF2BP3 expression level than the S-pattern (Fig. 3F, 3G).

Gene expression analysis using the public database The Cancer Genome Atlas revealed significantly higher IGF2BP3 levels in CRC tissue than in normal colon tissue (Fig. 3H), which is consistent with the findings of a previous report(Chen *et al*, 2023). In a database that included adenomas, although the sample size was small, the distribution of IGF2BP3 expression in adenomas was statistically more diverse than that in normal tissues, with higher expression levels in some adenomas (Fig. 3I). These results suggested that the *IGF2BP3* gene expression level increases during carcinogenesis.

### IGF2BP3 was functionally involved in the differences in D and S patterns

The functional role of IGF2BP3 in adenoma was also examined. First, *IGF2BP3* was knocked down in the D-pattern organoids (CP1-1, CP3-2, and CP9-2) (Fig. 4A). The D-pattern shifted to the S-pattern in the SSFG assay, and the SFC decreased in all cases when *IGF2BP3* was knocked down (Fig. 4B). Staining for cleaved caspase-3 revealed enhanced cell death (Fig. 4C and 4D). In contrast, Ki67 staining showed that proliferation was not significantly altered by IGF2BP3 knockdown (Fig. 4C, S3A). Therefore, IGF2BP3 knockdown promoted cell death and resulted in an S-pattern in D-pattern organoids. Knockdown of IGF2BP3 increased the protein level of CDK6 and the phosphorylation of S6, and clearly increased the expression level of MYC, suggesting that the cell cycle and cell biogenesis were activated (Fig. 4E). *IGF2BP3* was overexpressed in the S-pattern organoids (CP3-1, CP39-2, and CP1-5) (Fig. 4F). CP3-1 and CP1-5 shifted to the D-pattern in the SSFG assay when *IGF2BP3* was overexpressed, whereas CP39-2 maintained the S-pattern (Fig. 4G). SFC was higher when IGF2BP3 was overexpressed in all cases. Ki67 positivity increased, but that of cleaved caspase-3 did not change after overexpression of IGF2BP3 in all cases, including CP-39-2 (Fig. 4H, 4I, S3B). MYC protein levels and S6 phosphorylation levels increased (Fig. 4J). Taken together, these results indicated that IGF2BP3 plays different roles in S-pattern and D-pattern adenomas. In the S-pattern organoids, an increase in IGF2BP3 protein levels led to a transition to the D-pattern by promoting proliferation, whereas in D-pattern organoids, IGF2BP3 was involved in preventing cell death. MYC may play an important role as a downstream target of IGF2BP3, since protein levels are tightly related to IGF2BP3 levels, although in opposite directions. IGF2BP3 promotes MYC expression in S-pattern organoids, but suppresses it in D-pattern organoids.

**Figure 4.**
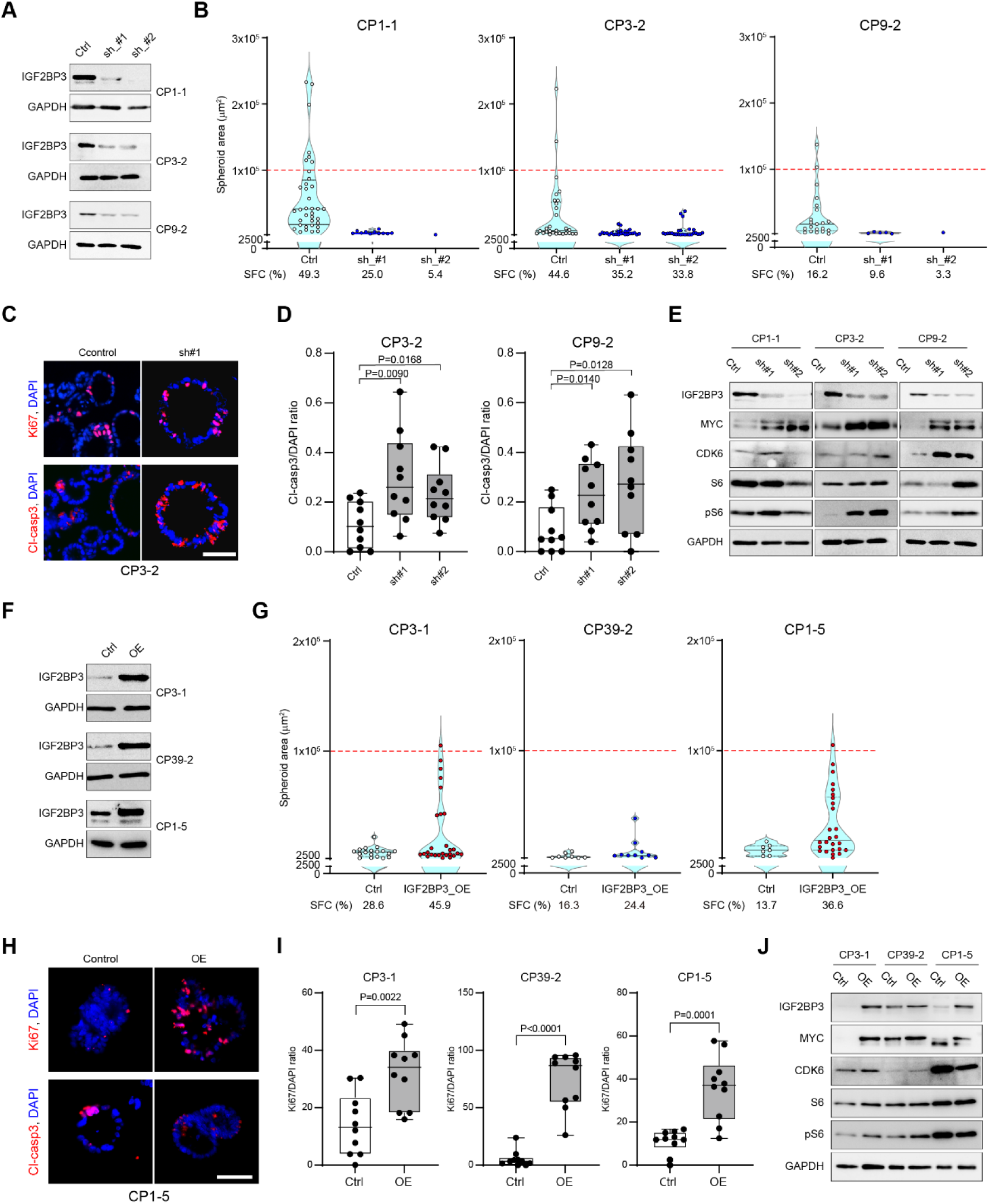
IGF2BP3 was functionally involved in the differences in D and S patterns **A**) Western blotting analyses of IGF2BP3 and the indicated proteins in D-pattern organoids infected with a lentivirus that expresses *IGF2BP3* shRNA#1 (sh#1), *IGF2BP3* shRNA (sh#2) or a scrambled shRNA (Ctrl). **B**) Violin plot of the SSFG assay for the indicated D-pattern organoids comparing control and IGF2BP3-knockdown cells. **C**) Immunofluorescence images of Ki-67 and cleaved caspase-3 in control and shIGF2BP3 organoids. **D**) Quantitative analysis of cleaved-caspase-3-positive ratios in (**C**). **E**) Western blotting analyses of the IGF2BP3, MYC, CDK6, S6, and pS6 in the indicated organoids. **F**) Western blotting analyses of IGF2BP3 and the indicated proteins in S-pattern organoids infected with a lentivirus that constitutively expresses IGF2BP3 (IGF2BP3_OE) or with the corresponding empty vector (Ctrl). **G**) Violin plot of the SSFG assay for the indicated S-pattern organoids comparing control and the IGF2BP3-overexpressing cells. **H**) Immunofluorescence images of Ki-67 and cleaved caspase-3 in control and IGF2BP3-overexpressing cells. **I**) Quantitative analysis of Ki67-positive ratios in (**H**). **J**) Western blotting analyses of IGF2BP3, MYC, CDK6, S6, and pS6 in the indicated organoids.

To identify the target genes of IGF2BP3, RIP sequencing was performed using D-pattern organoids. In the GO analysis, biological process terms related to the cell cycle were highly enriched (Fig. S4A), and hallmarks related to RAS and WNT signaling, as well as MYC targets, were also enriched (Fig. S4B). IGF2BP3 bound not only to *MYC* mRNA but also to many transcripts in the MYC signaling pathway, suggesting that IGF2BP3 is involved in regulating MYC signaling (Fig. S4C, S4D).

### IGF2BP3 regulated c-MYC at the transcriptional and translational levels

We investigated the relationship between IGF2BP3 and MYC expression. First, using MYC-specific RIP-qPCR, we confirmed that IGF2BP3 directly bound to *MYC* mRNA (Fig. 5A). IGF2BP3 positively regulates MYC by increasing the stability of *MYC* mRNA(Huang *et al*, 2018). In addition, when IGF2BP3 was overexpressed in CP1-5, an S-pattern organoid, *MYC* mRNA levels increased, but when it was knocked down in CP1-1, a D-pattern organoid, *MYC* mRNA levels decreased (Fig. 5B). In contrast, the protein level of MYC increased when IGF2BP3 was knocked down in D-pattern organoids (Fig. 4E). Thus, there is a discrepancy between the regulation of MYC mRNA and protein expression by IGF2BP3. A decrease in degradation is unlikely to be the cause of the increased protein levels, because the degradation rate of the MYC protein was almost the same as that of the control when IGF2BP3 was knocked down (Fig. 5C). In contrast, when IGF2BP3 was knocked down, the proportion of *MYC* mRNA in the polysome fraction increased (Fig. 5D, 5E). This suggested that IGF2BP3 negatively regulates MYC translation in the D-pattern organoids.

**Figure 5.**
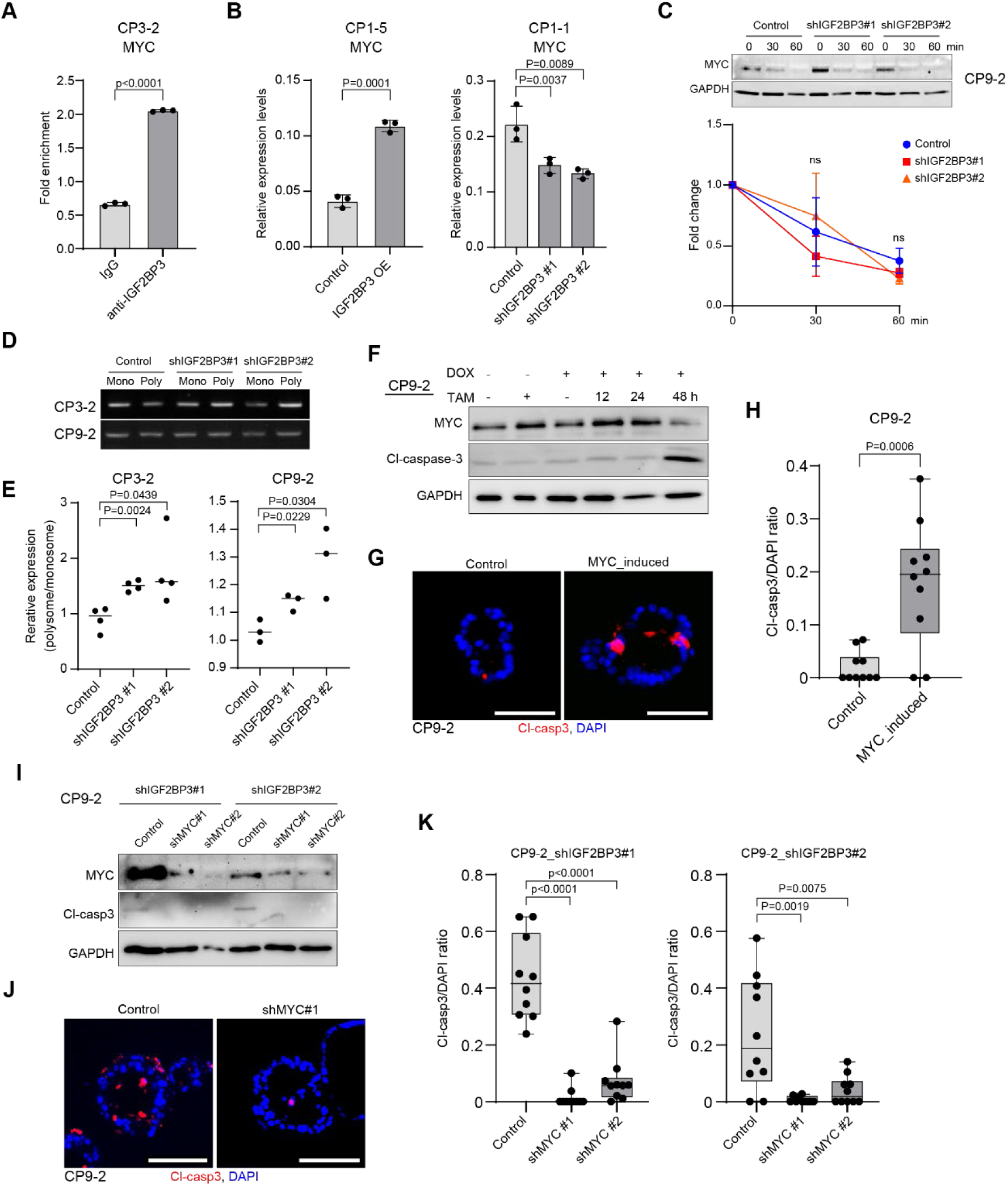
IGF2BP3 regulated c-MYC at the transcriptional and translational levels **A**) RNA immunoprecipitation (RIP)-quantitative polymerase chain (PCR) of the *MYC* gene using a negative control and anti-IGF2BP3 antibody. The ratio compared to each input is shown. **B**) Relative *MYC* gene expression levels in the indicated IGF2BP3-overexpression (left graph) and IGF2BP3-knockdown (right graph) organoids, assessed by reverse transcription (RT)-PCR. **C**) Cycloheximide (CHX) chase analysis of MYC protein. Time course of western blotting 0, 30, and 60 min after CHX treatment in D-pattern organoids (upper) and the quantitative analysis (lower). **D**) Polysome fractionation and detection of *MYC* mRNA. *MYC* mRNA was detected in monosome (mono) and polysome (poly) fractions using RT-PCR. **E**) *MYC* mRNA was quantified in (**D**) and the polysome/monosome ratio of *MYC* mRNA is shown. **F**) Western blotting of MYC and cleaved caspase-3 in D-pattern organoids with a doxycycline-inducible MYC-ER vector. Organoids were treated with tamoxifen and doxycycline as indicated. **G**) Immunofluorescence images of cleaved caspase-3 in control and MYC-induced organoids **H**) Quantitative analysis of the cleaved-caspase-3-positive ratio in (**G**). **I**) Western blotting of MYC and cleaved caspase-3 in the D-pattern organoid, CP9-2 with IGF2BP3 knockdown infected with lentiviruses expressing *MYC* shRNA#1 (shMYC#1), *MYC* shRNA#2 (shMYC#2), or a scrambled shRNA (Control). **J**) Immunofluorescence analysis of cleaved caspase-3 in control and shMYC#1 CP9-2 organoids. (**K**) Quantitative analysis of the cleaved-caspase-3-positive ratio in (**J**).

MYC overexpression in normal cells induces apoptosis(McMahon, 2014). When MYC was overexpressed in an inducible manner in D-pattern organoids, the MYC protein was induced, with a peak at 24 h, and cleaved caspase-3 increased after 48 h (Fig. 5F, 5G, 5H). MYC levels decreased at 48 h, likely because of the induction of the death of cells with high MYC expression levels. We further investigated whether cell death induced by IGF2BP3 knockdown could be rescued by MYC suppression. In cells with double knockdown of IGF2BP3 and MYC, the levels of cleaved caspase-3 were suppressed (Fig. 5I). Immunostaining revealed a significant decrease in the number of cleaved-caspase-3-positive cells (Fig. 5J, 5K). This suggested that cell death induced by IGF2BP3 knockdown was due to MYC overexpression. These results suggested that IGF2BP3 contributes to the maintenance of MYC expression at appropriate levels in D-pattern adenomas by promoting *MYC* transcription, while repressing translation to avoid cell death due to MYC overexpression.

### Overexpression of IGF2BP3 increased tumorigenicity *in vivo*

We previously investigated the tumorigenicity of mouse intestinal organoids in which driver gene mutations were introduced and reported that organoids with RNAi-mediated *Apc* knockdown were already tumorigenic(Onuma *et al*, 2013). We obtained similar results by knocking out *Apc* by introducing Cre via lentivirus infection *in vitro* in intestinal organoids prepared from Apc^flox/flox^ mice. In the meantime, we obtained a sub-cultured line by chance in which tumorigenicity was lost when the organoids were subcutaneously transplanted into mice (Fig. 6A; 6F, control). The subcultured line may have lost some of the elements necessary for tumor formation. The expression levels of Myc and Igf2bp3 diminished after long-term culture (Fig. 6B).

**Figure 6.**
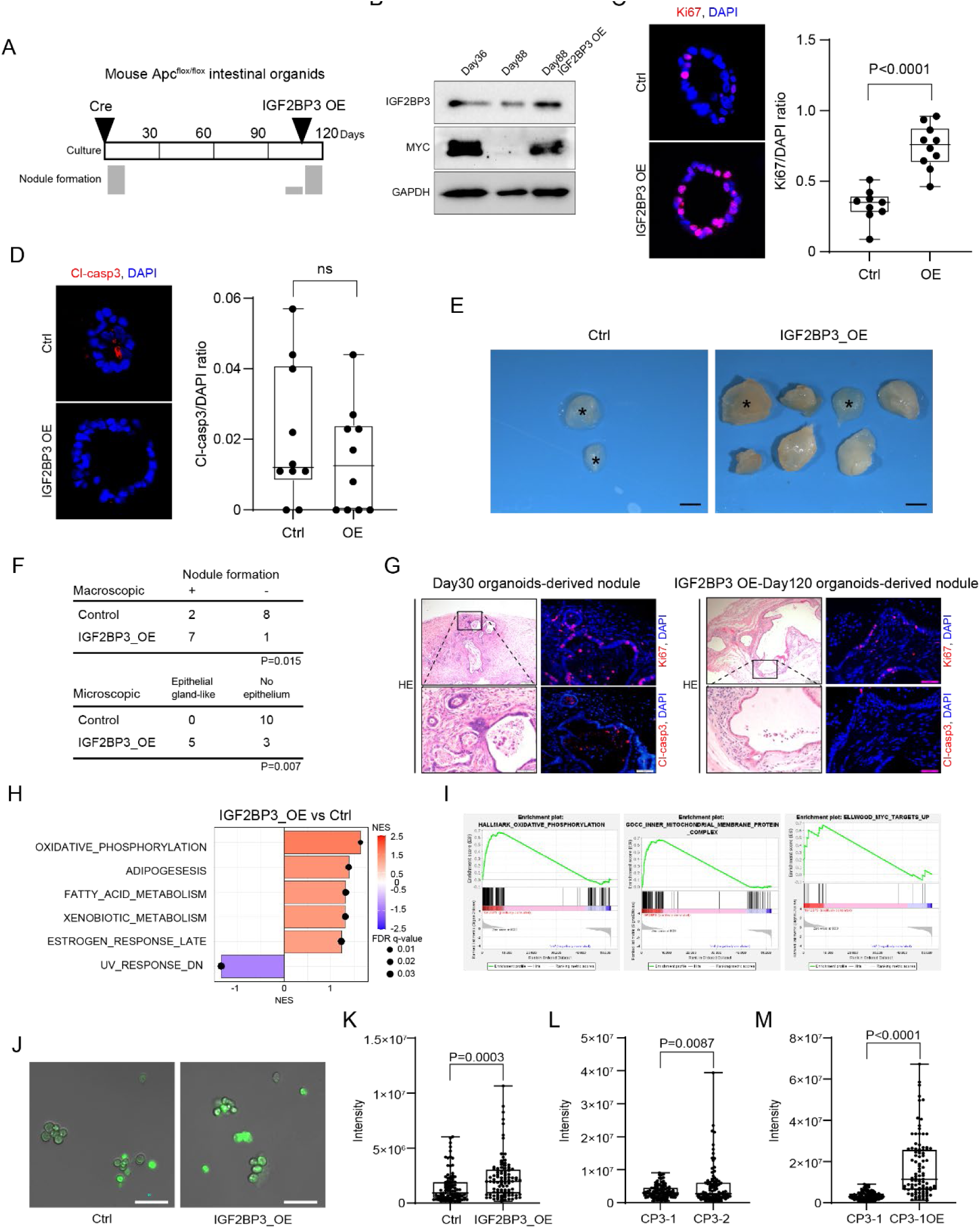
Overexpression of IGF2BP3 increased tumorigenicity *in vivo* **A**) Overview of the experimental schedule for *Apc* knockout, IGF2BP3 introduction, and tumorigenicity in NOD SCID mice. **B**) Western blotting analyses of Igf2bp3 and Myc in the indicated organoids. **C, D**) Immunofluorescence images of Ki-67 (**C**) and cleaved caspase-3 (**D**) in control and IGF2BP3-overexpressing cells. Ki-67- and cleaved-caspase-3-positive ratios were quantified. **E**) Macroscopic images of the excised nodules developed from injected organoids. Asterisks indicate nodules with no detectable epithelial component. Scale bar, 2 mm. **F**) Summary of nodule formation after inoculating the indicated organoids. **G**) Representative H&E and immunohistochemical images of Ki67 and cleaved caspase-3 (cl-casp3) in nodules derived from day 30 organoids and IGF2BP3-OE day 120 organoids. Scale bar, 50 μm. **H**) Hallmark pathway analysis comparing IGF2BP3_OE and control organoids. The bar graph represents the pathways, with the bubble size proportional to the false discovery rate (FDR) q-val and a color gradient indicating the normalized enrichment score (NES). Red bars denote pathways that are upregulated and positively enriched in IGF2BP3_OE organoids, while blue bars signify pathways that are downregulated and negatively enriched in IGF2BP3_OE organoids. **I**) GSEA of the transcriptome data using the indicated signatures comparing the control (Ctrl) and IGF2BP3_OE organoids. **J**) Representative microscopic images of the cells from organoids stained with MitoTracker Green. Green, MitoTracker; blue, Hoechst33342. Scale bar: 50 μm. **K**, **L**, **M**) Quantification of MitoTracker Green staining. The fluorescence intensity per cell is shown. Each dot represents a cell.

Although overexpression of IGF2BP3 did not promote tumorigenesis in human adenoma organoids, we examined whether overexpression of IGF2BP3 could reverse tumorigenesis in long-term-cultured *Apc*-knockout mouse intestinal organoids. As previously reported, the SSFG assay cannot be performed on mouse intestinal organoids because of their low SFC, as previously reported(Hall *et al*, 2022). IGF2BP3 protein levels increased with forced expression and Myc protein levels increased in parallel (Fig. 6B). Ki67 staining showed increased proliferation (Fig. 6C), whereas cleaved caspase-3 staining showed no change in cell death (Fig. 6D). These results are consistent with those of human S-pattern organoids. When organoids with IGF2BP3 overexpression (IGF2BP3-OE) were inoculated into immunodeficient mice, tumorigenicity was restored; however, microscopically, the nodules were epithelial-gland-like and not adenomatous (Fig. 6E, 6F). The histology of the IGF2BP3-OE nodules resembled that of the nodules at 30 days after *Apc* knockout (Fig. 6G). Ki67 and cleaved caspase-3 staining showed that cell proliferation and cell death were similar to those of the nodules on day 30 (Fig. 6G).

Gene expression profiling comparing long-term cultured *Apc*-knockout organoids with IGF2BP3-OE organoids revealed that the MYC signature was enriched upon the overexpression of IGF2BP3 (Fig. 6I, S5A). Oxidative phosphorylation and metabolic pathways were enriched in IGF2BP3-OE cells (Fig. 6I, S5B). These results are consistent with those of the microarray analysis of patient-derived adenoma organoids, in which these signals were enriched in the D-pattern organoids (Fig. 3). When mitochondrial activity in the cells was evaluated using MitoTracker Green, mouse IGF2BP3-OE showed more active mitochondria than the control cells (Fig. 6K). D-pattern human organoids showed higher mitochondrial activity than S-pattern organoids (Fig. 6L, S5C), and IGF2BP3 overexpression in S-pattern organoids resulted in increased mitochondrial activity (Fig. 6M). These results suggested that IGF2BP3 regulates MYC expression at an optimal level in mouse models, contributing to tumor formation in the adenoma-carcinoma sequence.

## Discussion

This study revealed that the intra-tumor heterogeneity of cell proliferation observed in cancer has already been observed in some adenomas. Acquisition of the D-pattern, that is, the epi-genetic and reversible promotion of proliferative potential in some cells, may be a step in the adenoma-carcinoma sequence. IGF2BP3 is expressed at a higher level in the D-pattern organoids than the S-pattern organoids. When IGF2BP3 is overexpressed, the S-pattern becomes a D-pattern, and when it is knocked down in the D-pattern organoids, the D-pattern phenotype is lost. Therefore, IGF2BP3 expression is an important factor in the formation of D-patterns in adenomas. IGF2BP3 may function as an epi-genetic step in the carcinogenic process by activating the cell cycle and preventing cell death by regulating MYC protein levels. There are both similarities and differences between D-pattern adenomas and cancers. Spheroids derived from the L cells of D-type adenomas gave rise to both S-and L-cells (Fig. 2A), similar to the situation in CRC(Coppo *et al*., 2023a). Thus, the D-pattern organoids are not solely composed of cells with a high proliferative potential. Although the possibility that the acquisition of the D-pattern is genetically determined cannot be ruled out, the proliferative range is not genetically controlled. The SFC was used to determine the characteristics of stem and progenitor cells. In CRC, there was no difference in SFC between the S-pattern and D-pattern(Coppo *et al*., 2023a). By contrast, in adenomas, the D-pattern had a higher SFC than the S-pattern (Fig. 1D), and the D-pattern may be closer to the stem-like state. In adenomas, we observed an exceptionally fast-growing spheroid in the first-round SSFG assay, which gave rise to S-pattern spheroids in the second round (Fig. 2A, #136). This exceptional phenomenon was not observed in CRC(Coppo *et al*., 2023a), indicating that the D-pattern phenotype is not as strict in adenomas. MYC expression was higher in the D-pattern both in cancer(Coppo *et al*., 2023a) and adenoma tissues (Fig. 2D, 2E). Therefore, MYC may play an important role in regulating cell proliferation. In contrast, IGF2BP3 was highly expressed in CRC cells (Fig. 3H), and there was no difference between the patterns(Coppo *et al*., 2023a). The expression level of MSI1, which plays an important role in the transition of cancer growth patterns(Coppo *et al*., 2023a), did not differ between the adenoma patterns. Therefore, there are similarities in the variations in growth as a phenotype between cancer and adenoma; however, there are significant differences in the underlying mechanisms.

IGF2BP3 is highly expressed in many types of cancers and has been reported to be a poor prognostic factor(Liu *et al*, 2023). IGF2BP3 knockdown inhibits the growth of CRC cell lines(Xu *et al*, 2019; Yang *et al*, 2020). The target genes of IGF2BP3 in cancer are diverse, and IGF2BP3 has been reported to stabilize mRNAs, including *EGFR*(Chen *et al*., 2023), *VEGF*, cyclin D1(Yang *et al*., 2020), *CDK6*(Palanichamy *et al*, 2016), and *HOXA*(Tran *et al*, 2022) mRNA, and promote their transcription. In addition to direct effects, it may also have indirect effects via the destabilization of p53 by stabilizing the regulatory factors of the signaling pathway(Zhao *et al*, 2017). In addition, IGF2BP3 has been reported to stabilize COX6B2 in lung cancer cells and activate oxidative phosphorylation, making the cells resistant to EGFR tyrosine kinase inhibitors(Lin *et al*, 2023), which is consistent with the fact that IGF2BP3 activates mitochondria in a D-pattern (Fig. 6L). Since RIP-seq analysis revealed that IGF2BP3 bound to the signaling molecules RAS and WNT (Fig. S4B), as well as MYC in a D-pattern organoid, IGF2BP3 may contribute to carcinogenesis in multiple ways.

MYC is highly expressed in CRC(Cancer Genome Atlas, 2012). In normal intestinal homeostasis, MYC plays a fundamental role in regulating cell proliferation and crypt cell number(Muncan *et al*, 2006). In contrast, acute MYC overexpression increased p53 and ARF levels and induced apoptosis. MYC also triggers apoptosis by inducing Bim, which neutralizes Bcl-2 (Dang *et al*, 2005). Thus, MYC upregulation occurs during the transition from a normal epithelium to an adenoma, and controlled upregulation of MYC can lead to cellular proliferation, thus avoiding cell death(Ben-David *et al*, 2014). MYC drives global metabolic reprogramming at the adenoma stage, regulates many metabolic reactions, and alters the expression levels of numerous metabolic genes(Satoh *et al*, 2017). However, the inter- and intra-tumor heterogeneity of adenomas has not been assessed in previous studies.

MYC protein levels were higher in D-pattern organoids than in S-pattern organoids (Fig. 2D, 2G), and cell death did not increase (Fig. S2A). The anti-death mechanism may have already been acquired in the D-pattern organoids, although the exogenous expression of MYC induced cell death in these organoids (Fig. 5F, 5G, 5H). The regulatory mechanisms controlling MYC expression levels within a narrow range in adenomas are largely unknown. The findings of this study suggest that the upregulation of IGF2BP3 expression may be responsible for the upregulation of MYC expression and threshold control in adenomas. IGF2BP3 binds to *MYC* mRNA to stabilize it, which increases its transcription level(Palanichamy *et al*., 2016). Indeed, *MYC* transcription was upregulated by IGF2BP3 overexpression in S-pattern organoids (Fig. 5B). Moreover, IGF2BP3 is reported to bind to *MYC* mRNA and induce its translocation from the polysome to the P-body, thereby inhibiting translation(Shan *et al*, 2023). Indeed, forced suppression of IGF2BP3 increased MYC protein levels in the D-pattern organoids (Fig. 4E). Thus, IGF2BP3 stimulated *MYC* transcription and simultaneously suppressed MYC translation. It is possible that dual control by IGF2BP3 contributes to the regulation of MYC at an appropriate level for adenoma growth.

We have shown that IGF2BP3 regulates MYC in adenomas; however, the roles of other downstream genes of IGF2BP3 and upstream factors that increase IGF2BP3 expression levels will be elucidated in the future. MYC regulates IGF2BP3(Bell *et al*, 2013; Du *et al*, 2022); thus, the existence of a positive feedback loop must also be considered. Since no gene mutations discriminating between the S-pattern and D-pattern were detected (Table S3), it is possible that epigenetic transcriptional regulation, such as methylation or chromatin remodeling, is involved. Through the analysis of the upstream regulators of IGF2BP3 expression and downstream m6A modifications, the epigenetic mechanisms in the adenoma-carcinoma sequence will be elucidated in future studies. Furthermore, because IGF2BP3 is expressed at low levels in normal tissues, the inhibition of IGF2BP3 may be a therapeutic strategy that can prevent the progression of adenomas to cancer by inducing apoptosis through MYC overexpression, while minimizing the effects on normal tissues.

## Methods

### Preparation and culture of tumor organoids

This study was approved by the Institutional Ethics Committee of Kyoto University (G1035-5, R1671-5) and all human samples were obtained with written informed consent from the patients. Fresh specimens of endoscopically or surgically resected colorectal adenomas were obtained and divided into two parts, one for pathological analysis and the other for culture (Table S1). Morphologically normal mucosa was obtained from resected specimens of patients with colon adenoma or cancer. The adenomas were divided into two parts: one for pathology and the other for culture. The adenomas and normal mucosa were mechanically minced. Without enzyme digestion, the fragments were embedded in Matrigel GFR (Corning, Corning, NY, USA) and cultured in untreated plates (IWAKI, Tokyo, Japan) in the respective media (Table S2). Organoids were passaged *in vitro* every 4–7 days. They were incubated in TripLE Express (Thermo Fisher Scientific, Waltham, MA) at 37℃ for 7 min, pipetted, and embedded in 5 µL of Matrigel GFR (Corning, 354230). All experiments were performed at least 24 h after passaging. Organoid samples were also stored at -80℃ in CS10 medium (STEMCELL Technologies, Vancouver, Canada).

### SSFG assay

The SSFG assay was performed as previously described(Coppo *et al*., 2023b). To prepare single cells, the organoids were treated with 0.25% trypsin-EDTA (Thermo Fisher Scientific) and DNase I (10 µg/mL) for 10 min at 37℃ and 1 min at room temperature, respectively. The cell suspension was gently pipetted 100 times and filtered through a 35 μm cell strainer (Corning) to remove cell clusters. The dissociated single cells were diluted in SSFG medium (adenoma organoid medium containing 2% Matrigel GFR and 10 μM ROCK inhibitor [Y-27632]; Selleck Chemicals, Houston, TX, USA), and seeded in a non-treated 384-well plate (Sumitomo Bakelite, Tokyo, Japan) at a ratio of 1 cell per well (50 mL/well). Within 2 h of cell seeding, microscopic images of each well were acquired using a LEICA DMI4000B microscope (Leica Microsystems, Wetzlar, Germany) equipped with an XY stage (Mitani Corporation, Tokyo, Japan). Wells containing strictly single cells with an area less than 300 µm^2^ on day 0 were used for subsequent analyses. Fresh SSFG medium (30 µL/well) was added on day 7. The area of each single-cell-derived spheroid was measured using the acquired microscopic images on day 14 and ImageJ Fiji software. Spheroids larger than 2,500 µm^2^ were defined as growing spheroids. The spheroid-forming capacity (SFC) was calculated as the percentage of the number of growing spheroids to the total number of single cells on day 0. The D-pattern was defined as the formation of at least one spheroid ≥ 1.0 × 10⁵ μm² at day 14, while the S-pattern was defined as the formation of spheroids exclusively < 1.0 × 10⁵ μm². This threshold was applied based on a previous report(Coppo *et al*., 2023a).

### Animal studies

Animal studies were approved by the Institutional Animal Care and Use Committees of Kyoto University (18564) and the Chiba Cancer Center (51) and were performed in compliance with institutional guidelines. For the tumor-formation assay, organoids equivalent to 1 × 10^6^ cells were suspended in a 1:1 mixture of medium and Matrigel (Corning) and injected into the flank of NOD/Scid mice (4–5 weeks old) (CLEA Japan, Tokyo, Japan). Apc^flox/flox^ mice(Shibata *et al*., 1997) were obtained from Ryoji Yao, the Japanese Foundation for Cancer Research. The classification of subcutaneous nodules was based on a previous report(Maru *et al*, 2019).

### RNA immunoprecipitation (RIP)-sequencing/RIP-quantitative reverse transcription polymerase chain reaction (qRT-PCR)

RIP was performed using a Magna RIP RNA-Binding Protein Immunoprecipitation Kit (EMD Millipore, Billerica, MA, USA) according to the manufacturer’s protocol. Sequence library preparation and next-generation sequencing were performed by Takara Bio, Inc. (Kusatsu, Japan). Briefly, the SMART-Seq v4 Ultra Low Input RNA Kit (Takara Bio) was used to prepare the sequencing library and 150-base pair-end sequence analysis was performed using a NovaSeq 6000 instrument (Illumina). RIP-qPCR was performed using an anti-human IGF2BP3 antibody (Proteintech, San Diego, CA, USA) or normal rabbit IgG, followed by qRT-PCR, as described below.

### Separation of polysomal RNA fractions

Polysome fractionation was performed as previously described(Endo *et al*, 2017; Okuyama *et al*, 2010). Briefly, cells were lysed in cell lysis buffer comprising 1% Triton X-100, 0.3 M NaCl, 15 mM MgCl_2_, 15 mM Tris-HCl (pH 7.4), and 100 U of RNase inhibitor (Promega, Madison, WI, USA) for 10 min on ice. The nuclear fraction and cell debris were removed by sequential centrifugation. The RNA content in the cytoplasmic extracts was determined and extracts containing equal amounts of RNA were overlaid on a 10-50% sucrose gradient. After ultracentrifugation in an Optima-XE100 (Beckman Coulter, Brea, CA, USA) at 36,000 rpm for 120 min at 4°C, the lysates were separated into monosomal or polysomal fractions and resolved by 1% agarose gel electrophoresis. RNA was extracted from each fraction using an RNeasy Mini Kit (QIAGEN). The quantity of *MYC* transcripts in each fraction was determined by semiquantitative RT-PCR using SuperScript II reverse transcriptase (Invitrogen, Carlsbad, CA, USA) and Takara Ex Taq (Takara Bio).

### Cycloheximide chase analysis

Cycloheximide chase analysis of MYC was performed as previously described (Okuyama *et al*., 2010). Organoids were treated with 50 μg/mL cycloheximide (Sigma-Aldrich), and proteins were extracted after 0, 30, and 60 min. MYC protein was detected using western blotting.

### *Apc* knockout in mouse intestinal organoids

Organoids from the small intestines of Apc^flox/flox^ mice(Shibata *et al*., 1997) were isolated and cultured as previously described(Onuma *et al*., 2013). *Apc* was deleted by lentiviral introduction of Cre recombinase using LV-Cre pLKO.1 (Addgene #25997), which encodes Cre recombinase in the backbone of the pLKO.1 vector. To select *Apc*-knockout cells, R-spondin1-depleted culture medium was used, as it does not allow the propagation of Apc^flox/flox^ cells without recombination. Successful recombination was confirmed by genotyping the transduced organoids for the *Apc* allele, as previously described(Onuma *et al*., 2013).

## Statistical analysis

Statistical analyses were performed using GraphPad Prism, version 9 (GraphPad Software). Significance was tested using the unpaired Student’s t-test for single comparisons and one-way or two-way analysis of variance followed by Tukey’s or Bonferroni’s test for multiple comparisons. For analyses of the SSFG assay results that did not show a normal distribution, a non-parametric Mann–Whitney test was used. Statistical significance was set at *p* < 0.05.

## Disclosures

### Ethics Statement

-Approval of the research protocol by an Institutional Reviewer Board: This study was approved by the Institutional Ethics Committees at Kyoto University (R1575, R1671).
- Informed Consent: Fresh biopsy specimens from patients were obtained after obtaining written informed consent.
- Animal Studies: All animal experiments were approved by the Institutional Animal Care and Use Committee of Kyoto University (18564). All procedures were performed in compliance with institutional guidelines.

### Transcript Profiling

GSE291602 https://www.ncbi.nlm.nih.gov/geo/query/acc.cgi?acc=GSE291602

GSE291756 https://www.ncbi.nlm.nih.gov/geo/query/acc.cgi?acc=GSE291756

GSE291843 https://www.ncbi.nlm.nih.gov/geo/query/acc.cgi?acc=GSE291843

## Writing assistance

Funding: This work was supported by the Japan-UK Research Cooperative Program between JSPS and the Royal Society (M.I.) and grant from JSPS KAKENHI, 22K07149 (C) (A.Y., M.I.).

## Author Contributions

T. S., Formal Analysis, Investigation, Methodology, Visualization, Writing – original draft; R. C., Formal Analysis, Writing – review & editing; K. O., Supervision, Writing – review & editing; J. K., Supervision, Writing – review & editing; H. U., Investigation, Writing – review & editing; D. H., Investigation, Resources,; Y. Y., Formal Analysis, Writing – review & editing, O. K., Supervision, Writing – review & editing; S. O., Supervision, Writing – review & editing; Y. T., Supervision, Writing – review & editing; K. K., Formal Analysis; Y. H., Supervision; Y. I., Resources, Writing – review & editing; K. O., Resources, Writing – review & editing; Y. H., Conceptualization, Supervision, Resources, Writing – review & editing; M. M., Supervision, Project administration, A. Y., Conceptualization, Funding acquisition, Supervision, Project administration, Writing – review & editing; M. I. Conceptualization, Funding acquisition, Supervision, Project administration, Visualization, Writing – original draft

## Abbreviations

CHX: cycloheximide
CRC: colorectal cancer
FAP: familial adenomatous polyposis
GO: Gene Ontology
GSEA: gene set enrichment analysis
IGF2BP3: insulin-like growth factor 2 mRNA-binding protein 3
KO: knockout
m6A: N6-methyladenosine
mRNA: messenger RNA
NOM: nominal
OE: overexpression
RIP-seq: RNA immunoprecipitation sequencing
qRT-PCR: quantitative reverse transcription polymerase chain reaction
SFC: spheroid-forming capacity
SSFG assay: single-cell-derived spheroid-forming and growth assay

